## Supplemental data for "LncRNA *Snhg3* Aggravates Hepatic Steatosis via PPARγ Signaling"

**table supplement 1. Liver lncRNA-Seq of DIO mice.**

**table supplement 2. Liver RNA-Seq between WT and *Snhg3*-HKI mice.**

**table supplement 3. Liver ATAC-Seq between WT and *Snhg3*-HKI mice.**

**table supplement 4. DARs associated with DEGs.**

**table supplement 5. Integrated analysis with ATAC-Seq and RNA-Seq.**

**table supplement 6. *Snhg3*-bound proteins identified by RNA-Pulldown-MS.**

**table supplement 7. *Snhg3*-bound proteins predicted by RBPsuite.**

**table supplement 8. Cut&Tag-Seq for H3K27me3.**

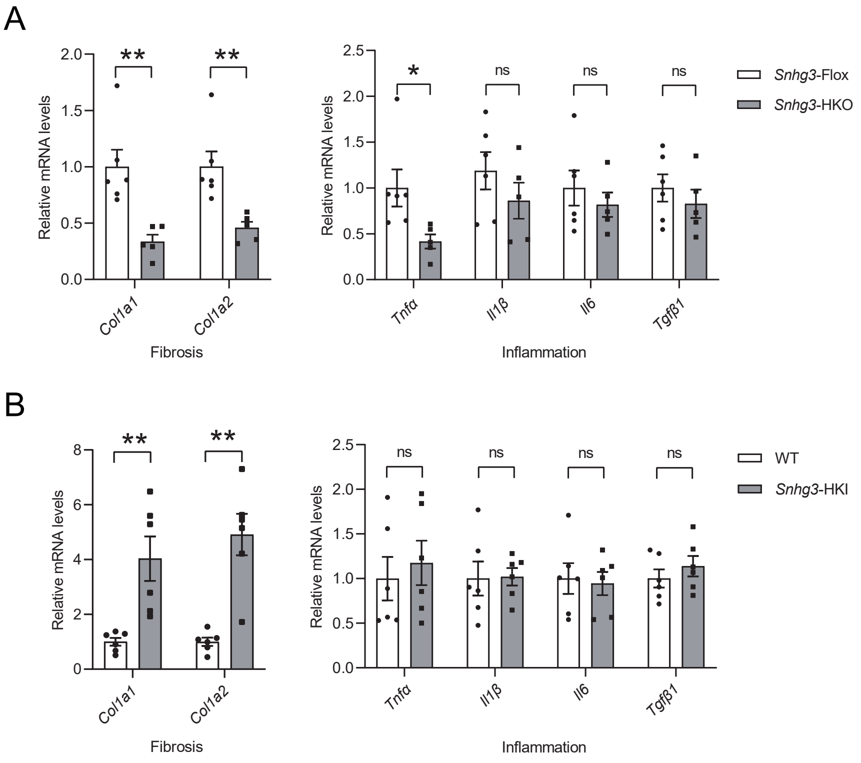


**figure supplement 1.** The mRNA levels of liver fibrosis and inflammation in DIO *Snhg3*-HKO mice (**A**) and *Snhg3*-HKI mice (**B**), compared to the controls. Data are represented as mean ± SEM. **$p < 0.01$ by Student's t test.

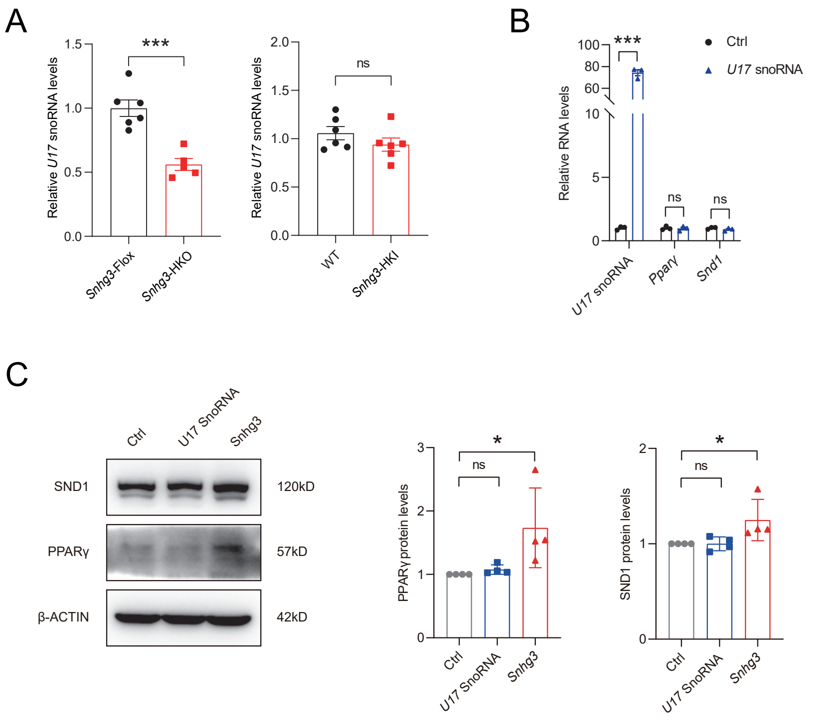


**figure supplement 2.** (**A**) Hepatic U17 snoRNA expression in DIO *Snhg3*-HKO mice and *Snhg3*-HKI mice compared to the controls. (**B** and **C**) Overexpression U17 snoRNA has no effect on the mRNA (**B**) and protein (**C**) levels of PPARγ and SND1 (left, western blotting; right, quantitative result). Data are represented as mean ± SEM. *$p < 0.05$ and ***$p < 0.001$ by Student's t test.
